# Healing of chromosomal breaks is impeded in cells expressing progerin

**DOI:** 10.64898/2026.08.13.744695

**Authors:** Aubrey A. Bondurant, Emma K. Grove, Nina M. Van, Alannah J. DiCintio, Alan S. Waldman

## Abstract

Hutchinson-Gilford Progeria Syndrome (HGPS) is a rare genetic condition characterized by features of accelerated aging, with a life expectancy of less than two decades. HGPS is commonly caused by a point mutation in the LMNA gene which codes for lamin A, a vital component of the nuclear lamina. The HGPS mutation activates a cryptic splice site and leads to production of a truncated, farnesylated form of lamin A referred to as "progerin." Progerin is also produced in small amounts in healthy individuals and has been implicated in normal aging. HGPS is associated with an accumulation of genomic DNA double-strand breaks (DSBs), and alterations in DSB repair. DSB repair in mammalian cells normally occurs by either homologous recombination (HR), an accurate, templated form of repair, or by DNA end-joining (EJ), a non-templated rejoining of DNA ends. EJ is error-prone, although a portion of EJ events occurs precisely with no alteration to joined sequences. Previously, we reported that over-expression of progerin increased EJ relative to HR and decreased the precision of EJ. In our current work, we designed a novel model experimental system using derivatives of thymidine kinase (tk)-deficient mouse fibroblasts and incorporating a loss-of-function assay to further explore progerin’s impact on EJ. We established cell lines containing an integrated copy of a functional herpes tk gene with an embedded recognition site for endonuclease I-SceI. We examined EJ at the nucleotide level following induction of a DSB within the tk gene by expression of I-SceI and subsequent selection for cells that lost tk gene function. Comparison of EJ products recovered from cells expressing progerin versus from cells not expressing progerin revealed that progerin expression provoked larger DNA deletions associated with DSB repair as well as recovery of multiple repair products from individual cells, suggesting progerin impedes re-joining of DNA ends.

## Introduction

Mammalian cells continually face a plethora of DNA damage, and maintenance of genome stability depends on the efficacy of a set of DNA repair pathways. One form of damage that cells must cope with is the DNA double-strand break (DSB). DSBs can be produced by chemical or radiological insult, and they may also form spontaneously from other DNA lesions or at stalled or collapsed replication forks. Quick and accurate repair of DSBs can help a cell avoid potentially deleterious genetic rearrangements and mutations.

To heal DSBs, mammalian cells have two general modes of repair at their disposal: homologous recombination (HR) and direct DNA end-joining (EJ) [reviewed in 1-10]. Although there are a few different types of HR events, and there are several mechanistically distinct pathways that may fall under the umbrella term EJ, the seminal difference between the repair strategies of HR versus EJ is that HR utilizes a template sequence which serves to maintain genetic information at the DSB site that may otherwise be altered, while EJ involves no template in the rejoining of DNA ends. Additionally, HR is active primarily during the late S or G2 stage of the cell cycle in dividing cells, whereas EJ is active throughout the cell cycle and in non-dividing cells. HR and EJ normally function in combination to preserve genome integrity, but corruption of either mode for repair may lead to harmful genomic change. For example, the potential exists for abnormally high levels of HR to lead to loss-of-heterozygosity of deleterious alleles, or the production of deletions or inversions between repeated sequences [11,12]. Further, mammalian cells normally allow HR exchange to occur only between sequences that exhibit a very high degree of sequence identity [13–15]. Genetic perturbations that allow the choice of inappropriate HR partners can lead to genomic instability in the form of localized sequence alterations, chromosomal rearrangements, or translocations [10,16,17]. When it comes to EJ, the lack of a repair template may produce sequence deletions or insertions, although a significant portion of end-joining events proceeds precisely with no alteration to the sequences at the break site [18–22]. We refer to precise end-joining as PEJ, and imprecise end-joining, in which sequences are altered as a consequence of repair, as IEJ. The degree to which a cell uses PEJ versus IEJ, the size of any sequence deletions or insertions associated with IEJ, as well as the appropriate regulation of HR, are all pertinent to the maintenance of genome stability. Of course, the overall ability of a cell to heal a DSB by any means is also crucial to genome stability.

The consequences of loss of genome stability can prove detrimental in a number of ways. It is well-documented that aberrant DNA repair is often associated with cancer [23–25]. Improper genome maintenance has also been implicated in aging. A slew of reports documents an increase in genomic instability that accompanies aging, correlating with an alteration of the intrinsic efficiency or nature of a variety of DNA repair pathways [26–39]. Alterations in EJ were reported in rat brain during aging [32], and studies with mice have suggested that the fidelity of DSB repair diminishes with age [33]. Both the efficiency and fidelity of EJ has been observed to decrease as human fibroblasts approach senescence [34]. Chromosomal DSBs accumulate in human cells approaching senescence, and it has been suggested that DSBs may be involved directly in the induction of senescence [26]. In short, evidence abounds for a role for impaired or altered repair of DSBs in the biology of aging. It is not difficult to appreciate that an age-associated decrease in genome integrity, along with possible apoptotic responses to unrepaired DNA lesions, may compromise critical cellular functions.

Genetic disorders that produce clinical features of premature aging are often associated with DNA repair defects and genomic instability [27–31, 35–39]. Hutchinson-Gilford Progeria Syndrome (HGPS) is one such rare genetic syndrome that leads to accelerated aging [40]. The average lifespan of an individual with HGPS is about fourteen years. The most common cause of HGPS is a point mutation in the LMNA gene which normally codes for lamin A and its splice variant lamin C. The mutation responsible for HGPS activates a cryptic splice site which leads to the production of a truncated form of lamin A referred to as "progerin." Interestingly, it has been learned that progerin is in fact expressed at low levels in healthy individuals and appears to play a role in the normal aging process [41–44]. Unlike wild-type fully processed lamin A, progerin retains a farnesyl and a methyl group at its carboxy terminus. The farnesyl and methyl groups cause progerin to largely remain associated with the inner nuclear membrane rather than localize to the nuclear lamina where lamin A normally resides.

Lamin A is an important component of the nuclear lamina, a structure that resides just inside of the inner nuclear membrane and is comprised of a fibrous meshwork of intermediate filaments and associated proteins. The nuclear lamina plays structural and catalytic roles in the nucleus [reviewed in 45]. In HGPS, the impact of progerin expression on nuclear architecture is profound. The nuclei of HGPS cells are characteristically misshapen and blebbed, and this altered nuclear structure conveys changes to numerous nuclear functions. Progerin expression in HGPS interferes with recruitment of replication factors to replication forks, leading to replication fork stalling and collapse [39,44–49]. Lamin A and its variants have also been directly implicated in impacting DNA repair [50–56].

Progerin expression in HGPS cells leads to an accumulation of DSBs and sensitivity to DNA damaging agents [45–49,57,58]. DSB repair in HGPS is apparently delayed, or precluded, due to delayed recruitment of DSB repair proteins, particularly those involved in HR repair, to sites of chromosomal damage in HGPS cells [46,58]. This delay in recruitment of HR-associated proteins is in line with observations suggesting that EJ is enhanced while HR is concomitantly reduced by progerin expression [52,53,55,59]. We previously confirmed this progerin-associated alteration in DSB repair by directly demonstrating that progerin expression shifts repair pathway choice at a defined genomic DSB away from the HR pathway and towards EJ [60]. We more recently used a novel model cell culture system using mouse fibroblasts to show that progerin expression alters the very nature of EJ by bringing about a marked shift away from PEJ and toward IEJ [61]. Our data also suggested that progerin expression correlates with an increase in deletion size associated with IEJ [61].

In our previous studies on the impact of progerin on DSB repair [60,61], DSB repair events were recovered using a model system in which we selected for the gain-of-function of a selectable marker gene following DSB induction. Such an approach necessarily restricts the types of events that can be retrieved. We have now designed an experimental system that uses a loss-of-function selection to recover cells following IEJ at a defined DSB. This strategy allows the recovery of a more comprehensive spectrum of DSB-induced events. Using this newly developed system, we show that progerin-expression does not appear to diminish a cell’s ability to heal a DSB by IEJ per se. However, we confirm that the size of deletions produced at a DSB is increased in cells expressing progerin and we show that progerin expression is coupled with an increase in events that appear to have proceeded in multiple steps, or that result in the production of multiple IEJ products. Collectively, our results reveal a progerin-associated complexity in the manner by which DNA ends are joined, suggesting an impediment to EJ in cells expressing progerin.

## Materials and Methods

### General cell culture

All cell lines were derived from Ltk^-^ mouse fibroblasts and were grown in Dulbecco’s Modified Eagle Medium (DMEM/low glucose) supplemented with 10% heat-inactivated fetal bovine serum, minimal essential medium non-essential amino acids, and gentamicin (50 μg/ ml). Cells were maintained in a 37°C incubator in a 5% CO_2_ atmosphere.

### DSB repair reporter cell lines

A cell line that enables recovery of DSB repair events using a loss-of-function selection was engineered in multiple steps, beginning with a cell line containing an integrated copy of plasmid pTKSce2. pTKSce2, described previously [18], is based on the vector pJS-1 [62,63], which is a derivative of pSV2neo [64]. pTKSce2 contains a nonfunctional copy of the herpes simplex virus type one thymidine kinase (tk) gene on a 2.5-kb BamHI fragment inserted into the unique BamHI site of the vector. The tk gene was rendered nonfunctional by insertion of a 47 bp oligonucleotide after position 963 of the tk gene coding region (numbering as described by Wagner et al. [65]). The 47 bp oligonucleotide contains two 18 bp recognition sites for endonuclease I-SceI, and pTKSce2 was designed as a reporter of both PEJ as well as IEJ. Previously, substrate pTKSce2 had been transfected into mouse Ltk-fibroblasts and a cell line designated line “13” containing a single integrated copy of pTKSce2 was isolated as described [61]. A schematic of the nonfunctional tk gene in pTKSce2 is presented at the top of Fig 1A.

**Fig 1.**
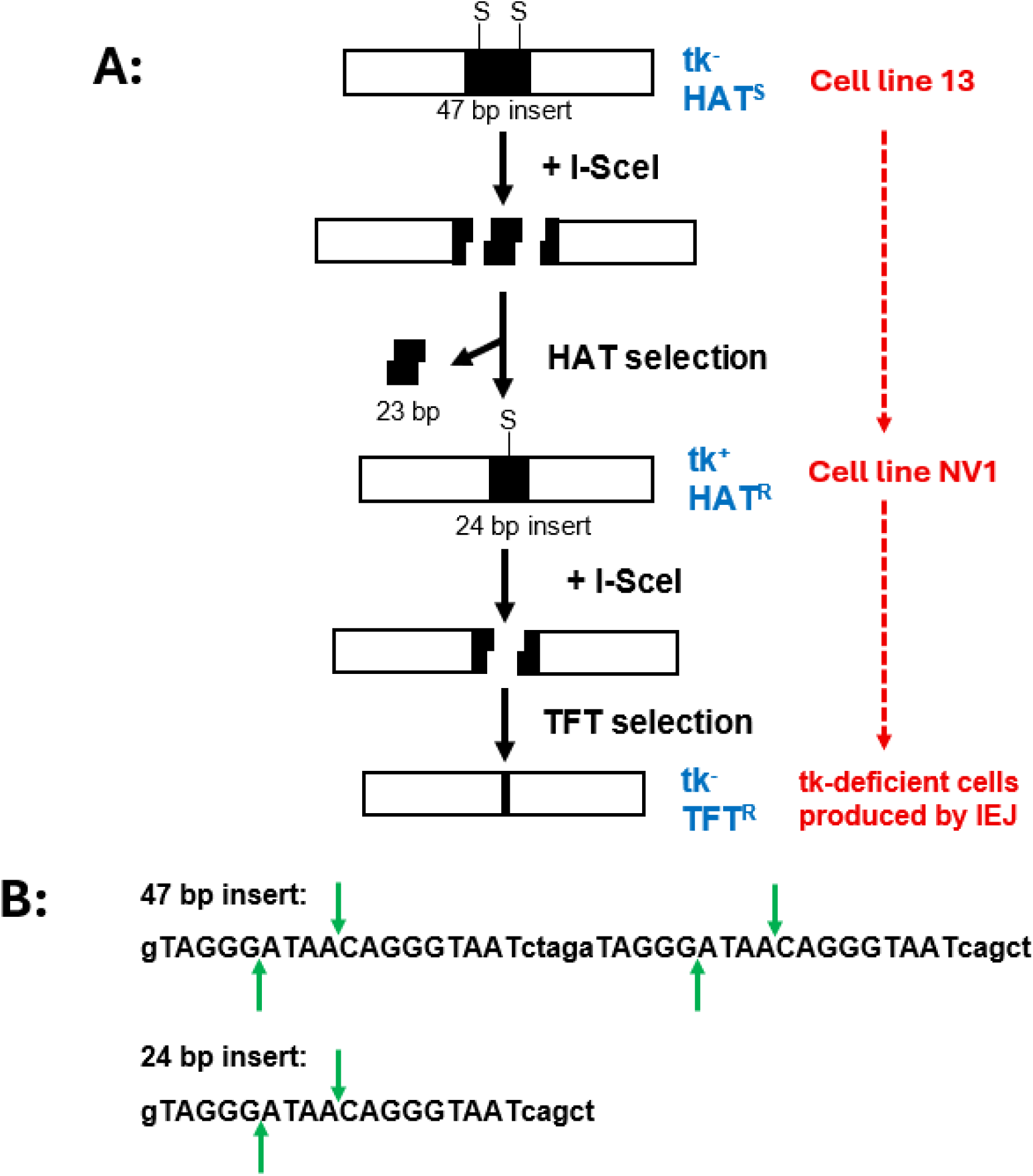
An experimental scheme to report IEJ using loss-of-function selection. (A) At the top of the figure is a schematic of the nonfunctional herpes tk gene in substrate pTKSce2, which was stably transfected into mouse Ltk^-^ cells to produce cell line13. The tk gene is disrupted by a 47 bp insert containing two I-SceI sites (labeled “S”). Subsequent cleavage at the two I-SceI sites followed by precise repair of the outer sticky ends produced cell line NV1. Cell line NV1 contains a functional tk gene harboring a single I-SceI site within a 24 bp insert. Cleavage at the I-SceI site within the tk gene in NV1 followed by IEJ at the DSB can render the tk gene nonfunctional, and the consequent tk-deficient cells can be selected in TFT. (B) Illustrated are the 47 bp and 24 bp inserts contained in the tk genes in cell lines 13 and NV1. The 18 bp I-SceI recognition sequence is in uppercase font, with green arrows indicating the staggered sites of I-SceI cleavage.

Cell line 13 was electroporated with I-SceI expression plasmid pCMV-3xnls-I-SceI (“pSce”) [66] to induce DSBs at the I-SceI sites within pTKSce2. Briefly, 5 × 10^6^ cells were mixed with 20 μg of pSce in a volume of 800 μl of phosphate buffered saline in a cuvette with a 0.4 cm electrode gap at room temperature and electroporated in a Bio-Rad Gene Pulser (Bio-Rad, Hercules, CA, USA) set to 750 V and 25 μFd. Following electroporation, cells were plated into hypoxanthine/aminopterin/thymidine (HAT) medium [67] to select for tk^+^ cells as described [61]. Among the tk^+^ colonies recovered, we identified a clone designated “NV1” that had undergone PEJ between the two outermost complementary DNA overhangs produced at the two I-Sce-I DSBs, as illustrated in Fig 1. NV1 was subsequently used to study DSB with a loss-of-function selection.

To produce derivatives of cell line NV1 that stably express GFP-progerin, 5 × 10^6^ NV1 cells were electroporated as above with a mixture of 10 μg of plasmid pEGFP-D50 lamin A (Addgene plasmid #17653) that had been linearized with *Eag*I, and 1 μg of pBABE-puro (Addgene plasmid #1764) which contains a puromycin resistance gene. Stably transfected clones were selected in puromycin (5 μg/ml). After 14 days of selection, colonies that showed nuclear GFP fluorescence were propagated further and GFP-progerin expression was confirmed by Western blot.

### Recovery of DSB repair events using loss-of-function selection

Prior to DSB induction, cell line NV1 and derivatives of NV1 expressing GFP-progerin were cultured in HAT medium for 4 days to kill any cells that may have spontaneously mutated to a tk-deficient phenotype. HAT selection was removed, and cells were allowed to continue to grow in DMEM for 5 additional days before DSB induction. DSBs were induced by electroporation of 3 x 10^5^ cells of a particular cell line were resuspended in 300μl of PBS containing 7.5 μg of pSce. Electroporation conditions were 750 V and 25 μFd using a cuvette with a 0.4 cm electrode gap. For mock experiments, cells were electroporated in PBS alone. After electroporation, cells were plated at a density of 1.25 x 10^4^ cells per 75-cm^2^ flask. The cells were grown in DMEM for 6 days under no selection and then refed with medium supplemented with 5 μg of trifluorothymidine (TFT)/ml to select for tk-deficient cells. Cells were refed with TFT-containing medium every 3 days for 1 week until TFT-resistant (TFT^R^) colonies were picked. Individual TFT^R^ colonies were propagated further, and genomic DNA was prepared from the cells. Two electroporations with pSce and two mock electroporations were carried out for each cell line studied and results from the two electroporations were pooled.

### PCR amplification and DNA sequence analysis

A fragment of the tk gene spanning the original location of the I-SceI recognition site was amplified from 500 ng of genomic DNA isolated from TFT^R^ colonies using primers AW100 (5′-TAATACGACTCACTATAGGGTTGCGCCCTCGCCGGCAGC-3′) and AW133 (5′-CAGGAAACAGCTATGACCCGGTGGGGTATCGACAGAGT-3′). AW100 is composed of nucleotides 600–618 of the coding sequence of the HSV-1 tk gene (numbering according to [65]), with a T7 forward universal primer appended to the 5′ end of the primer. AW133 is composed of nucleotides 1786–1767 of the HSV-1 tk gene with an M13 reverse universal primer appended to the 5′ end of the primer. PCR was accomplished using Ready-To-Go PCR beads (Cytiva) and a “touchdown” protocol as previously described [68]. One μl of each 25 μl PCR reaction was displayed on an agarose gel to assess whether a product had been generated. PCR products were then sequenced using a T7 primer by Eton Bioscience, Inc. (Research Triangle Park, NC).

### Western blots

Blots were performed using SDS-PAGE with 6% stacking gels and 8% separating gels. Each lane contained 30 μg of total cellular protein, and Bio-Rad Precision Plus Kaleidoscope Protein Standard (#161–0375, 5 μl) was used for molecular weight markers. Proteins were transferred to a nitrocellulose membrane. Ponceau S staining was performed to check transfer efficiency and to confirm equal protein loading across lanes, and membranes were blocked in 2% non-fat milk in PBST (PBS plus 0.1% Tween 20) for two hours. The primary antibody used was GFP (B-2): sc-9996 (mouse monoclonal, from Santa Cruz Biotechnology, Inc.) at a dilution of 1:500 in 5% non-fat milk-PBST and incubated at 4 degrees Celsius overnight on a rocker. Primary antibody was removed after washing 3x with PBST for 10 min. Secondary antibody used was goat anti-mouse IgG-HRP: sc-2005 (Santa Cruz Biotechnology, Inc.) at a dilution of 1:1000 in 5% non-fat milk-PBST and incubated at room temperature for two hours. The membrane was washed 3x with PBST prior to being developed. Detection was accomplished using GE Healthcare Amersham ECL Select Western Blotting Detection Reagent.

## Results

### An experimental scheme for studying imprecise DNA end-joining events in mammalian cells

To study a broad spectrum of IEJ events in a mammalian genome, we derived cell line NV1 from mouse Ltk^-^ fibroblasts. NV1 cells contain a functional tk gene harboring an inserted 24 bp sequence which includes a recognition site for endonuclease I-SceI (Fig 1). Following electroporation of cell line NV1 with pSce to induce a DSB within the functional tk gene, cells that have lost tk function can be recovered by selection in TFT. Since there are many types of IEJ events that can lead to loss of tk gene function, cell line NV1 provides a useful means for retrieving a broad array of IEJ events. The experimental scheme is presented in Fig 1, and further details are provided under Materials and Methods.

We were interested in learning how expression of progerin may influence the manner in which a DSB is repaired. To do so, we transfected cell line NV1 with pEGFP-D50 lamin A and recovered derivatives of NV1 that stably express GFP-progerin. Three of these derivatives were named NV1-prog2, NV1-prog5, and NV1-prog15. Each of these NV1 derivative cell lines displays nuclear fluorescence due to expression of GFP-progerin (Fig 2A), and expression of GFP-progerin in each of these lines was confirmed by western blot (Fig 2B). The fluorescence images and western blot indicate that the level of expression of GFP-progerin is similar among lines NV1-prog2, NV1-prog5, and NV1-prog15.

**Fig 2.**
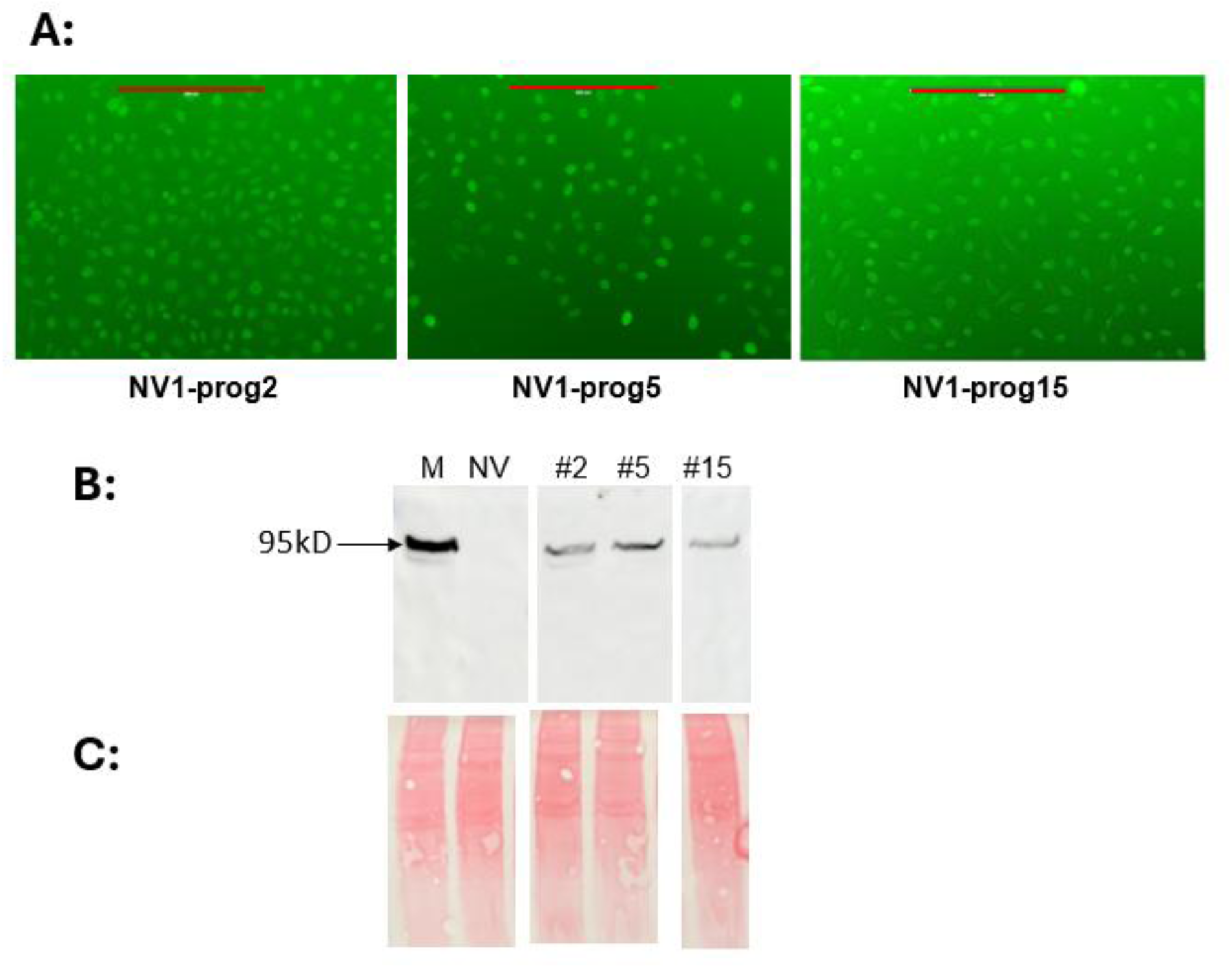
**Expression of GFP-progerin in derivatives of cell line NV1. (**A) Fluorescent microscopy images of cell lines NV1-prog2, NV1-prog5, and NV1-prog15 demonstrating nuclear fluorescence, indicative of GFP-progerin expression. The red bar in each image represents 200 microns. (B) Western blot using a GFP-specific antibody confirming expression of full-length GFP-progerin in cell lines NV1-prog2, NV1-prog5, and NV1-prog15 (lanes #2, #5, #15 respectively). Lane M displays a sample from murine cell line “alpha” that we had previously shown to express GFP-progerin at a level similar to but somewhat higher than progerin levels detected in cells derived from an HGPS patient [60,61], and this lane serves as a marker for GFP-progerin. Parent cell line NV1 (lane NV) does not display a band. Each lane contains 30 μg of total cellular protein. (C) Equal protein loading on the western blot is demonstrated by Ponceau staining of the membrane.

Following electroporation of cell line NV1 and its progerin-expressing derivatives with pSce and subsequent selection for tk^-^ colonies in TFT, comparison of the IEJ repair events recovered from the various lines allows an assessment of progerin’s impact on the nature of DSB repair.

### Cells expressing progerin are proficient at DSB repair via IEJ

Cells from line NVI and progerin-expressing derivative lines NV1-prog2, NV1-prog5, and NV1-prog15 were electroporated with pSce, or with PBS alone in mock electroporations, and placed under selection in TFT to recover tk-deficient cells following the regimen described under Materials and Methods. The numbers of TFT^R^ colonies recovered for each cell line are presented in Table 1. For each cell line, electroporation with pSce generated TFT^R^ colonies at a frequency more than an order of magnitude greater than the colony frequency seen following mock electroporation, indicating that almost all colonies recovered following electroporation with pSce were induced by a DSB. Also, the fact that the frequency of TFT^R^ colonies recovered from mock electroporations of progerin-expressing cells was not greater than observed for mock electroporations of parent cell line NV1 suggested that spontaneous mutation rate was not increased by progerin.

**Table 1.**
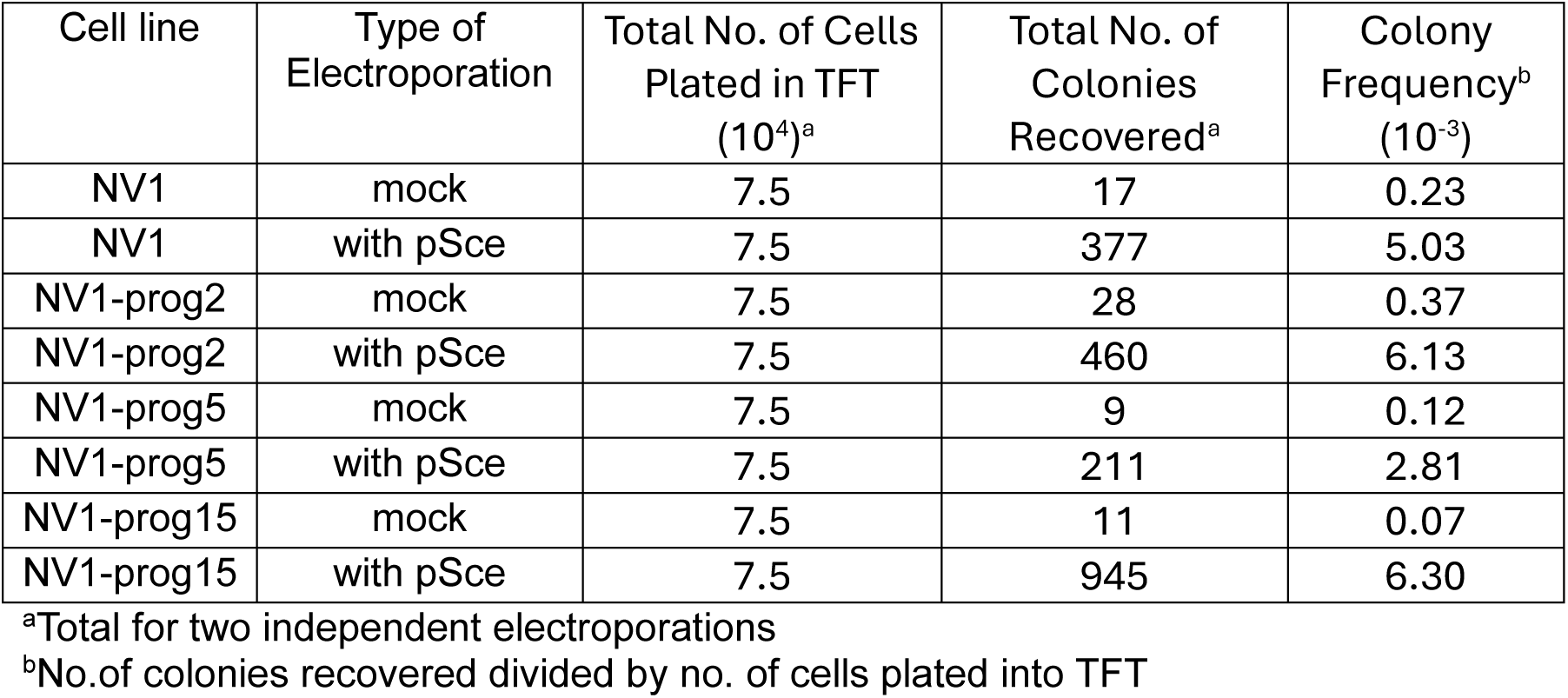
Recovery of DSB repair events.

| Cell line | Type of Electroporation | Total No. of Cells Plated in TFT (10<sup>4</sup>)<sup>a</sup> | Total No. of Colonies Recovered<sup>a</sup> | Colony Frequency<sup>b</sup> (10<sup>-3</sup>) |
| --- | --- | --- | --- | --- |
| NV1 | mock | 7.5 | 17 | 0.23 |
| NV1 | with pSce | 7.5 | 377 | 5.03 |
| NV1-prog2 | mock | 7.5 | 28 | 0.37 |
| NV1-prog2 | with pSce | 7.5 | 460 | 6.13 |
| NV1-prog5 | mock | 7.5 | 9 | 0.12 |
| NV1-prog5 | with pSce | 7.5 | 211 | 2.81 |
| NV1-prog15 | mock | 7.5 | 11 | 0.07 |
| NV1-prog15 | with pSce | 7.5 | 945 | 6.30 |

<sup>a</sup>Total for two independent electroporations
<sup>b</sup>No. of colonies recovered divided by no. of cells plated into TFT

For the three cell lines NV1-prog2, NV1-prog5, and NV1-prog15, the overall frequency of TFT^R^ colonies recovered following electroporation with pSce was 5.08 x 10^-3^, which is nearly identical to the colony frequency of 5.03 x 10^-3^ recorded for parent line NV1. The similar recovery of DSB-induced colonies in NV1 and its progerin-expressing derivatives suggested that cells expressing progerin remain competent at IEJ repair per se. Additional evidence that expression of progerin did not lessen a cell’s ultimate ability to carry out IEJ was demonstrated by PCR of the region of DNA surrounding the I-SceI site following DSB induction. Genomic DNA was isolated from each of the recovered TFT^R^ colonies, and PCR was carried out using primers AW 100 and AW 133 which flank the I-SceI site and are positioned about 1.2 kb apart. Production of a PCR product required rejoining of DNA ends initially generated at the I-SceI site. If processing of the DNA ends by deletion or addition of nucleotides were to occur prior to end-joining, successful PCR amplification could still transpire as long as the sites of primer annealing were not destroyed and remained linked, with a spacing sufficiently close together to allow DNA synthesis to extend from one primer to the other. Gross chromosomal rearrangements or extensive deletions or insertions would preclude successful PCR.

Genomic DNA isolated from all 37 TFT^R^ colonies analyzed that were recovered from NV1, and genomic DNA from 51 out of 53 TFT^R^ colonies recovered from NV1-prog2, NV1-prog5, and NV1-prog15 proved to be suitable templates for producing PCR products, indicating that joining of DNA ends induced at the I-SceI site occurred in nearly every colony recovered. Two out of 16 colonies analyzed from NV1-prog5 failed to produce a PCR product. Representative PCR products are displayed on the gels in Fig 3. PCR was also attempted on genomic DNA from four TFT^R^ colonies recovered from mock electroporations of NV1 and from eight TFT^R^ colonies recovered from mock electroporations of NV1-prog2. None of the colonies from mock electroporations produced a PCR product, perhaps indicating loss of the integrated reporter substrate in such colonies.

**Fig 3.**
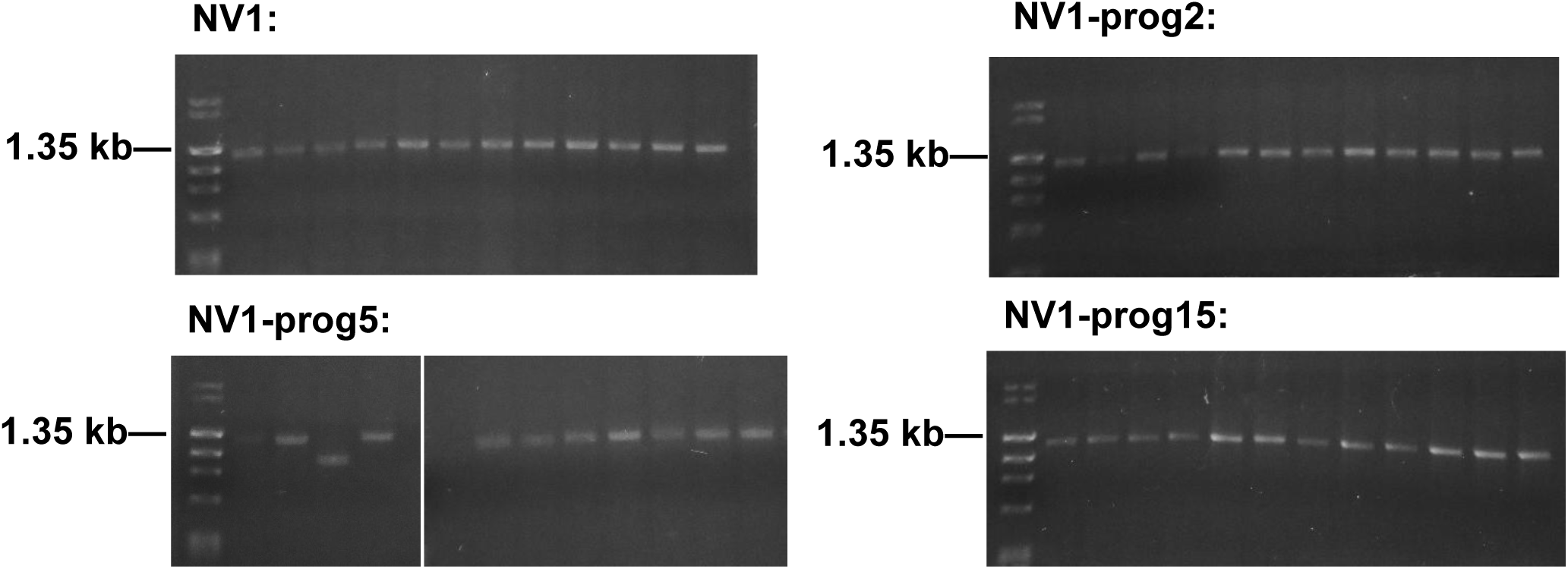
Representative PCR products generated from TFT^R^ colonies. Genomic DNA was isolated from TFT^R^ colonies following electroporation of cell lines NV1, NV1-prog2, NV1-prog5, and NV1-prog15 with pSce, and the DNA was used in PCR reactions with primers AW100 and AW133. The primers flank the I-SceI DSB site and are positioned about 1.2 kb apart prior to DSB-induction. Representative PCR products are displayed on agarose gels. The first lane of each gel contains DNA size markers, with the 1.35 kb band of a PhiX174 Hae III digest denoted. All samples analyzed from NV1 and almost all samples from the progerin-expressing derivatives generated PCR products.

### Nucleotide-level analysis of IEJ repair events suggests obstruction of repair in progerin-expressing cells

PCR products generated from TFT^R^ colonies were sequenced to further analyze the nature of IEJ repair in NV1 and its progerin-expressing derivatives, and the results are summarized in Tables 2-5. Sequence analysis revealed a higher degree of complexity among repair events recovered from the progerin-expressing lines. Compared with parent NV1, there were strikingly more colonies recovered from progerin expressing lines that contained a mixture of tk sequences. For NV1, 3 of 37 colonies analyzed revealed a mixture of two or more sequences, while 15 out of 53 colonies recovered from cells expressing progerin contained a mixture of sequences. This difference is significant (p = 0.0184 by chi square). Further, three TFT^R^ colonies from progerin-expressing cells had apparently undergone multi-stepped events. Colony #11 recovered from NV1-prog2 (Table 3) displayed a 3 bp deletion in conjunction with the addition of a T nucleotide, colony #13 from NV1-prog2 had two non-contiguous 1 bp deletions, and colony #14 from NV1-prog 15 (Table 5) displayed a 3 bp deletion in conjunction with the addition of an A nucleotide. When one considers colonies that either contained a mixture of sequences or had undergone a multi-stepped repair event, progerin-expressing cells produced highly significantly more such colonies than did NV1 (p = 0.0043). We also noted that 4 out of 37 IEJ events recovered from NV1 (Table 2) displayed a single A nucleotide inserted at the I-SceI site of strand cleavage, immediately following the ATAA sequence (see Fig 1), as the sole sequence change. In contrast, none of the 53 IEJ events recovered from progerin-expressing derivatives of NV1 (Tables 3-5) displayed a simple single nucleotide insert, again marking a significant difference compared with the parent NV1 line (p= 0.0143).

**Table 2.**
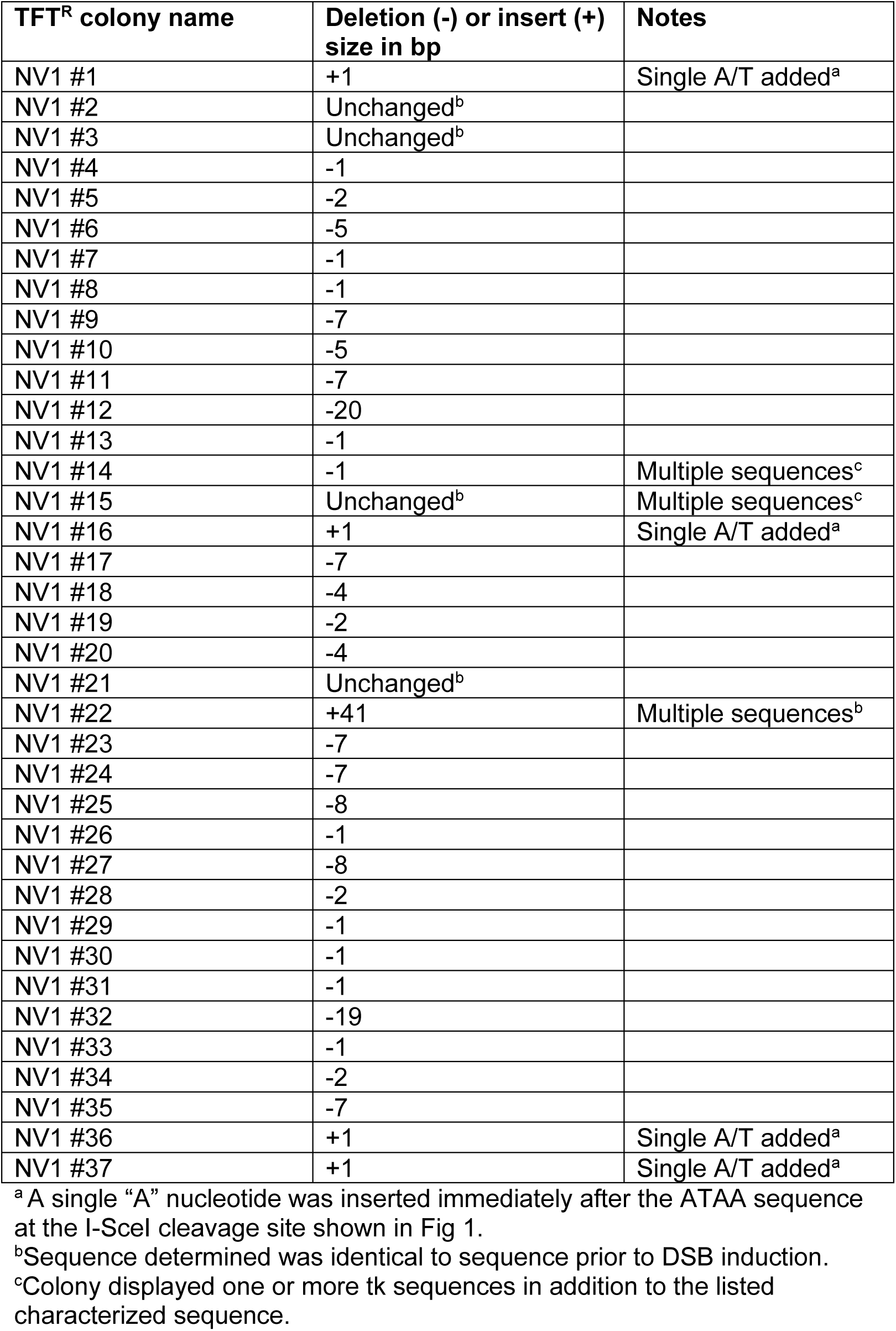
IEJ events recovered from cell line NV1.

| **TFT<sup>R</sup> colony name** | **Deletion (-) or insert (+) size in bp** | **Notes** |
| --- | --- | --- |
| NV1 #1 | +1 | Single A/T added<sup>a</sup> |
| NV1 #2 | Unchanged<sup>b</sup> |  |
| NV1 #3 | Unchanged<sup>b</sup> |  |
| NV1 #4 | -1 |  |
| NV1 #5 | -2 |  |
| NV1 #6 | -5 |  |
| NV1 #7 | -1 |  |
| NV1 #8 | -1 |  |
| NV1 #9 | -7 |  |
| NV1 #10 | -5 |  |
| NV1 #11 | -7 |  |
| NV1 #12 | -20 |  |
| NV1 #13 | -1 |  |
| NV1 #14 | -1 | Multiple sequences<sup>c</sup> |
| NV1 #15 | Unchanged<sup>b</sup> | Multiple sequences<sup>c</sup> |
| NV1 #16 | +1 | Single A/T added<sup>a</sup> |
| NV1 #17 | -7 |  |
| NV1 #18 | -4 |  |
| NV1 #19 | -2 |  |
| NV1 #20 | -4 |  |
| NV1 #21 | Unchanged<sup>b</sup> |  |
| NV1 #22 | +41 | Multiple sequences<sup>b</sup> |
| NV1 #23 | -7 |  |
| NV1 #24 | -7 |  |
| NV1 #25 | -8 |  |
| NV1 #26 | -1 |  |
| NV1 #27 | -8 |  |
| NV1 #28 | -2 |  |
| NV1 #29 | -1 |  |
| NV1 #30 | -1 |  |
| NV1 #31 | -1 |  |
| NV1 #32 | -19 |  |
| NV1 #33 | -1 |  |
| NV1 #34 | -2 |  |
| NV1 #35 | -7 |  |
| NV1 #36 | +1 | Single A/T added<sup>a</sup> |
| NV1 #37 | +1 | Single A/T added<sup>a</sup> |

<sup>a</sup> A single "A" nucleotide was inserted immediately after the ATAA sequence at the I-SceI cleavage site shown in Fig 1.
<sup>b</sup>Sequence determined was identical to sequence prior to DSB induction.
<sup>c</sup>Colony displayed one or more tk sequences in addition to the listed characterized sequence.

**Table 3.** IEJ events recovered from cell line NV1-prog2.

| **TFT<sup>R</sup> colony name** | **Deletion (-) or insert (+) size in bp** | **Notes** |
| --- | --- | --- |
| NV1-prog2 #1 | -1 |  |
| NV1-prog2 #2 | -11 |  |
| NV1-prog2 #3 | -2 |  |
| NV1-prog2 #4 | -1 |  |
| NV1-prog2 #5 | -23 |  |
| NV1-prog2 #6 | Complex | Mixture of several undetermined sequences |
| NV1-prog2 #7 | -5 | Mixture of sequences<sup>a</sup> |
| NV1-prog2 #8 | -2 |  |
| NV1-prog2 #9 | +257 |  |
| NV1-prog2 #10 | -40 |  |
| NV1-prog2 #11 | -3 | 3 bp deletion plus the addition of a T |
| NV1-prog2 #12 | +113 |  |
| NV1-prog2 #13 | -1, -1 | Two noncontiguous 1 bp deletions |
| NV1-prog2 #14 | mix |  |
| NV1-prog2 #15 | +5 |  |
| NV1-prog2 #16 | mix |  |
| NV1-prog2 #17 | -1 plus -1 | Mixture of two distinct deleted sequences |
| NV1-prog2 #18 | -13 |  |
| NV1-prog2 #19 | -5 |  |
| NV1-prog2 #20 | -5 |  |
| NV1-prog2 #21 | -5 |  |
| NV1-prog2 #22 | -2 |  |
| NV1-prog2 #23 | -29 |  |
| NV1-prog2 #24 | -4 |  |

<sup>a</sup>Colony displayed one or more tk sequences in addition to the listed characterized sequence.

**Table 4.** IEJ events recovered from cell line NV1-prog5.

| **TFT<sup>R</sup> colony name** | **Deletion (-) or insert (+) size in bp** | **Notes** |
| --- | --- | --- |
| NV1-prog5 #2 | -2 |  |
| NV1-prog5 #3 | -19 |  |
| NV1-prog5 #4 | -4 |  |
| NV1-prog5 #5 | -7 |  |
| NV1-prog5 #6 | -11 |  |
| NV1-prog5 #7 | -4 |  |
| NV1-prog5 #8 | -1 |  |
| NV1-prog5 #9 | -7 |  |
| NV1-prog5 #11 | -1 |  |
| NV1-prog5 #12 | -2 |  |
| NV1-prog5 #13 | complex | Mixture of several undetermined sequences |
| NV1-prog5 #14 | -20 |  |
| NV1-prog5 #15 | -285 |  |
| NV1-prog5 #16 | complex | Mixture of several undetermined sequences |

**Table 5.**
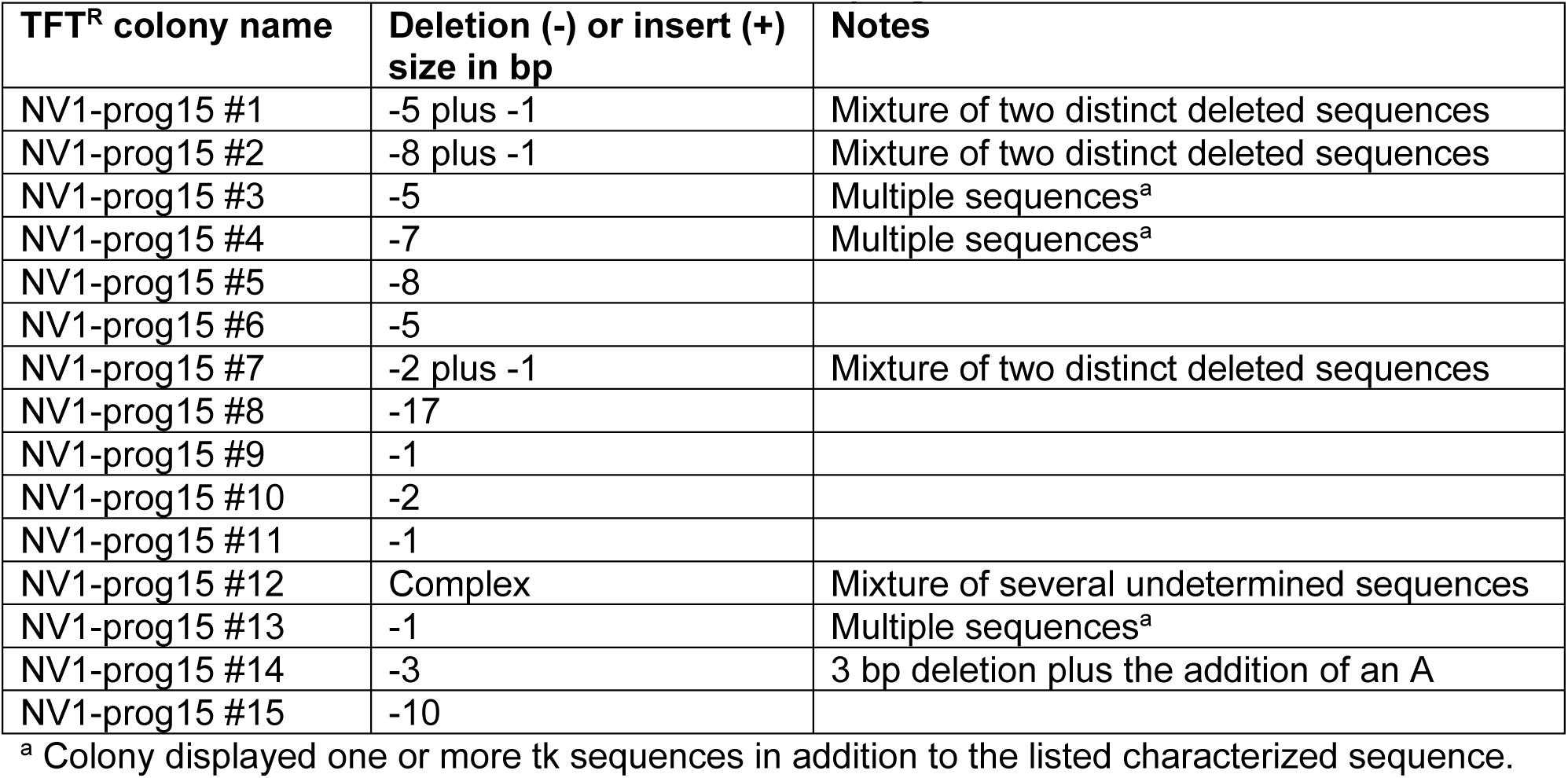
IEJ events recovered from cell line NV1-prog15.

| **TFT<sup>R</sup> colony name** | **Deletion (-) or insert (+) size in bp** | **Notes** |
| --- | --- | --- |
| NV1-prog15 #1 | -5 plus -1 | Mixture of two distinct deleted sequences |
| NV1-prog15 #2 | -8 plus -1 | Mixture of two distinct deleted sequences |
| NV1-prog15 #3 | -5 | Multiple sequences<sup>a</sup> |
| NV1-prog15 #4 | -7 | Multiple sequences<sup>a</sup> |
| NV1-prog15 #5 | -8 |  |
| NV1-prog15 #6 | -5 |  |
| NV1-prog15 #7 | -2 plus -1 | Mixture of two distinct deleted sequences |
| NV1-prog15 #8 | -17 |  |
| NV1-prog15 #9 | -1 |  |
| NV1-prog15 #10 | -2 |  |
| NV1-prog15 #11 | -1 |  |
| NV1-prog15 #12 | Complex | Mixture of several undetermined sequences |
| NV1-prog15 #13 | -1 | Multiple sequences<sup>a</sup> |
| NV1-prog15 #14 | -3 | 3 bp deletion plus the addition of an A |
| NV1-prog15 #15 | -10 |  |

<sup>a</sup> Colony displayed one or more tk sequences in addition to the listed characterized sequence.

Deletion sizes for IEJ events recovered from cells expressing progerin were larger than deletions associated with IEJ in parent NV1. Deletions of 10 bp or more were produced in two of the 37 IEJ colonies from NV1 and in 11 of 53 IEJ colonies from the progerin-expressing lines, a significant difference (p = 0.0415). Overall, the average size IEJ-associated deletion recovered from NV1 was 5.25 bp, while the average size deletion recovered from the progerin-expressing derivatives was 12.83 bp. Further, if one considers IEJ events that resulted in simply the deletion or addition of a single nucleotide at the DSB site,14 such events were recovered among the 37 events from NV1 whereas only 7 such events were among the 53 events recovered from cells that expressed progerin, a highly significant difference (p = 0.0066). Curiously, we also recovered sequences from 4 colonies from NV1 that were unchanged from the starting tk sequence.

As discussed below, our results are collectively consistent with an impediment to DSB repair in cells that express progerin.

## Discussion

In this work, we describe an experimental system using a loss-of-function selection to study the impact of progerin expression on DSB repair via IEJ in mammalian cells. Using our system, we recovered intrachromosomal IEJ events from murine fibroblast cell line NV1 and directly compared these repair events with IEJ events recovered from three progerin-expressing derivatives of NV1. This work complements our previous studies [60,61] in which we used a gain-of-function selection to study the difference in DSB repair events occurring in cells expressing progerin versus events occurring in cells that do not express progerin. The use of a negative selection in the current work allows a less restrictive recovery of IEJ events but necessarily precludes recovery of PEJ events.

In our earlier studies using a gain-of-function assay to recover DSB repair events from cell line 13 (Fig 1) and progerin-expressing derivatives of line 13, we reported a two- to three-fold lower frequency of recovery of repair events in the presence of progerin as well as a significant shift away from PEJ and toward IEJ [61]. We inferred that expression of progerin suppresses the annealing of terminal complementary bases, leading to a reduced ability of cells to successfully join DNA ends. In hindsight, if one considered only the IEJ events recovered in our earlier work [61], no deficiency in recovery of IEJ was evident in progerin-expressing cells relative to cells not expressing progerin. Cell line NV1 used in our current study was derived directly from line 13 (Fig 1), and our current work utilizes a loss-of-function assay to selectively recover *only* IEJ events at the very same genomic locus we investigated previously. Using line NV1, we report here that progerin expression does not reduce recovery of IEJ events. We conclude that while expression of progerin reduces a cell’s ability to carry out PEJ, it does not reduce a cell’s ultimate DNA end-joining capability.

Although progerin-expressing cells appear competent at joining DNA ends via IEJ in some fashion, sequence analysis revealed an impact of progerin on the nature of IEJ. Overall, IEJ events recovered from lines NV1-prog2, NV1-prog5, and NV1-prog15 were more complex than events from NV1. Significantly more colonies recovered from the progerin-expressing lines contained multiple sequences or had undergone multi-step events. Deletion size also was larger for events occurring in the presence of progerin, as we reported previously using a positive selection [61]. In the presence of progerin, significantly fewer IEJ events were recovered in which only a single nucleotide was either deleted or added at the site of strand incision. These observations, taken together, are consistent with the notion that DSB repair is impeded, and DNA ends are longer-lived prior to repair in cells expressing progerin. A longer life for DNA ends prior to repair in progerin-expressing cells would consequently provide time for increased nucleolytic degradation and/or increased opportunities for ends to participate in multiple processing steps prior to the ultimate joining of ends. The notion that progerin engenders longer-lived DSBs is in accord with previous reports of elevated levels of endogenous DNA damage and slow recruitment of repair proteins, mainly those involved in HR, to sites of DSBs in cells expressing progerin [45–49,58,69,70–73]. Our work suggests that this slowness of repair extends to EJ and is not restricted to HR.

A precise explanation for the elevated levels of accumulated damage seen in the presence of progerin has presently not been firmly established. We have reported [74] that much of this damage is not readily attributable to stalled replication forks, contrary to a popular paradigm [39,46–49,75–78]. We speculated [74] that despite the elevated levels of DNA damage seen with progerin, mitotically dividing cells may play “catch-up” when it comes to damage repair and are faced with the task of repairing elevated levels of damage in the course of a cell cycle. However, the burden of having to execute repair at an excess number of genomic sites may stretch a cell’s resources, including not only the availability of repair proteins but the availability of nucleotides as well. Cells expressing progerin may thus be “living on the edge” when it comes nucleotide pools. Progerin-associated accumulation of DNA damage coupled to inadequate nucleotide pools was implicated in our previous work [74] and has been demonstrated by others [79]. Limiting resources may prolong the life of a DSB even if it is eventually repaired.

How might multiple repair products be recovered from TFT^R^ colonies following IEJ, as seen in lines NV1-prog2, NV1-prog5, and NV1-prog15? It is possible that the copy number of the tk gene within some cells is amplified to two or more copies and that the multiple copies are repaired independently. If this were the case, then a Southern blot may reveal an amplified tk gene signal when genomic DNA is hybridized with a tk-specific probe. Southern blot analysis of representative colonies from which multiple tk sequences were recovered did not provide consistent convincing evidence for sequence amplification (data not shown). However, amplified sequence copy number might not be stable and thus might not always be so easily detected on a blot, and so it remains a possibility that at least some of the TFT^R^ colonies had undergone sequence amplifications.

A different, intriguing possibility for how a colony may contain multiple DSB repair products would be if the two DNA strands had been repaired independently and differently from one another. There is indeed evidence for independent, non-simultaneous repair of the two DNA strands at a DSB, at least under certain circumstances [80]. Normally, the second strand to be repaired at a DSB uses the initially repaired strand as a template, so the two strands of a repaired DNA duplex are typically fully complementary. However, should the two strands be repaired differently, mismatched heteroduplex would exist at least transiently. If the heteroduplex were to persist up to and through S-phase, a mixture of sequences would be produced by replication and would be recovered in a colony. It is thus possible that progerin expression leads to uncoupled repair of DNA strands. It is even imaginable that one strand is repaired while the other remains unrepaired up until replication. This may wreak havoc during replication, possibly leading to sequence amplification, and perhaps leading to some of the complex mixtures of sequences we recovered.

It is also formally possible that multiple, independent IEJ events may occur over a period of several days after plating cells following electroporation with pSce, and prior to killing by TFT selection. This would require persistence of I-SceI activity over several generations, with delayed DSB induction and repair after several days of growth. We do not consider this scenario to be very likely since we have not seen evidence of anything like this in our numerous previous studies involving DSB induction with I-SceI. More to the point, if simply delayed breakage and repair occurring over a period of days were the explanation for the recovery of mixed events in NV1-prog2, NV1-prog5, and NV1-prog15, we would expect to see a similar level of such occurrences in NV1, which we do not. The elevated level of mixed events in the progerin-expressing lines thus points to an impact of progerin and an associated generation of multiple repair products stemming from an initial single DSB.

The curious recovery of unchanged tk sequences, most notably from NV1, might be due to mutation outside of the bounds of the PCR products sequenced. It is also at least conceivable that recovery of unchanged tk sequences from tk-deficient colonies could be due to DSB-induced epigenetic silencing of the tk gene in combination with repair via PEJ. DSB repair is known to be often associated with localized transcriptional silencing that aids in timely repair and is normally transient in nature [81].

There is little disagreement that faltering pathways for genome maintenance constitute one aspect of the complex biology of aging. It is difficult to know whether any given change in DNA repair is a cause or an effect of aging, or if change in DNA repair and damage accumulation is actually better described as part of a viscous cycle of corruption of cellular processes that define the aging process. In any case, the more we catalog and learn about changes in cellular processes that accompany aging, the better equipped we will be to devise strategies for slowing aging or improving the quality of life as we age. Expression of progerin has not only been identified as the cause of the premature aging syndrome HGPS, it has also been implicated in normal aging as well, and so studying the impact of progerin expression on the nature of DNA repair can help us to make inroads into a more complete understanding of both accelerated and normal aging. Our current work adds to the broad picture that progerin expression interferes with efficient, accurate repair of chromosomal DSBs, which in turn may lead to genomic instability and eventual cellular and organismal demise. Defining the nature of a problem is a critical step in finding a solution. Armed with an expanding picture of the types of alterations in nucleic acid metabolism that aging may prompt, the challenging task of mitigating these changes awaits.

